# In vitro evolution of DNA operators enables multivalency protein−DNA interactions: towards programmable transcription factor regulation

**DOI:** 10.64898/2026.05.02.721436

**Authors:** Natalia S. Adler, Giuliano T. Antelo, Matías Villarruel Dujovne, Johnma J. Rondón, My T Le, David P. Giedroc, Ana Sol Peinetti, Daiana A. Capdevila

**Author notes:** To whom correspondence should be addressed (A.S.P.); (D.A.C.).

## Abstract

*In vitro* transcription (IVT) systems regulated by allosteric transcription factors (aTFs) are central to emerging cell-free biosensing and synthetic biology platforms, yet their performance is often limited by suboptimal protein–DNA interactions and the need for well-characterized regulatory elements. Here, we report an *in vitro* evolution strategy to engineer DNA operator sequences that enables tunable aTF–DNA interactions without requiring prior detailed knowledge of the native operator or regulatory mechanism. Using a SELEX-based approach with integrated positive and counter-selection steps, we evolved non-natural operators for the sulfane sulfur–responsive transcriptional repressor SqrR. The selected sequences preserve ligand-responsive allostery, with some sequences exhibiting enhanced binding affinity and reducing transcriptional leakage. Notably, we identify operator with binding behaviors consistent with cooperative recruitment of multiple SqrR dimers, suggesting that sequence architecture can modulate aTF–DNA interactions beyond affinity alone. Incorporation of these operators into IVT circuits improves transcriptional control and dynamic range, enabling the development of ROSALIND-based sensors for sulfane sulfur species, achieving sensitive and selective detection in a fully cell-free format. More broadly, this work establishes operator evolution as a programmable strategy to optimize transcription factor–DNA interactions and expand the design space of transcription-based biosensors, including for systems lacking well-characterized genetic components.

## INTRODUCTION

*In vitro* transcription (IVT) reactions constitute a versatile and powerful platform in synthetic biology and biotechnology, enabling the controlled production of RNA under defined conditions. IVT-based systems regulated by allosteric transcription factors (aTFs) that activate or repress downstream gene expression have shown great promise in diverse applications, including cell-free gene expression^1^ and the development of molecular diagnostics and biosensors.^2,3^ Transcription-based biosensing platforms such as ROSALIND have further demonstrated the potential of these systems for rapid and programmable molecular detection. In these reactions, aTFs bind specific DNA operator sequences to repress transcription and dissociate upon recognition of a cognate small-molecule ligand, thereby modulating transcription.

The performance of IVT-based regulatory systems critically depends on the strength and specificity of the interaction between the aTF and its cognate DNA operator. Efficient transcriptional control requires tight binding in the absence of the target ligand to effectively repress transcription, while remaining dissociable upon ligand binding at relevant analyte concentrations. Insufficient aTF–DNA affinity can result in transcriptional leakage, reduced signal-to-background ratios, and diminished amplification efficiency, ultimately compromising sensitivity and reliability. Likewise, systems where DNA affinity is too high and have low allosteric coupling lead to poor limits of detection (LoD). Therefore, engineering the aTF– operator interaction is essential to achieve optimal performance, but remains challenging due to the need to simultaneously balance multiple parameters.^4^ Some have addressed this challenge through high-throughput screening and directed evolution of aTFs, including approaches that combine cell-free expression platforms with automated screening and machine learning–guided optimization.^5,6^ More recently, others have sought to improve LoDs by either incorporating gene circuits that amplify the readout signal,^7^ or more sensible methods of detection that do not rely on fluorescence, such as lateral flow assays.^8^ While promising, these strategies rely on having a well characterized system in which both the regulatory mechanism of the aTF and its cognate DNA sequence is known.

While binding affinity is a central parameter governing transcriptional regulation, it does not fully capture the complexity of protein–DNA interactions. Increasing evidence indicates that the architecture of transcription factor binding sites, including their spacing, orientation, and multiplicity, can profoundly influence regulatory outcomes.^9,10^ In particular, the presence of multiple binding sites can enable cooperative recruitment of transcription factors, mediated by protein–protein interactions and DNA-dependent effects, and is recognized as an important determinant of gene regulatory behavior.^11,12^ Despite their relevance in natural gene regulation, these architectural features remain largely underexplored in the design of in vitro transcriptional circuits and cell-free biosensors.

To address this limitation, the *in vitro* selection of DNA sequences through the Systematic Evolution of Ligands by EXponential enrichment (SELEX) offers a powerful approach to identify operators with tailored functional properties. Originally developed to isolate high-affinity nucleic acid ligands, high-throughput SELEX approaches^13–15^ have been widely used to determine preferred binding motifs of DNA-binding proteins and to map the binding specificities of hundreds of human transcription factors.^16–18^ Biophysical models derived from SELEX-seq data have further helped elucidate how transcription factor orientation and spacing preferences influence protein–DNA interactions. Indeed, functional assays on the selected sequences obtained through SELEX allowed to assess how the subtle variations in operator length, symmetry, and nucleotide composition influence the stability and geometry of the protein–DNA complex, and affect transcriptional repression and inducibility. Consequently, operator optimization through SELEX provides a flexible route to fine-tune IVT regulatory circuits based on any aTF of interest. However, their application to the functional optimization of operator sequences in IVT-based systems has not been yet explored. Here, we designed an *in vitro* selection strategy incorporating both positive and counter-selection steps to obtain non-natural DNA operators that retain analyte-responsive allosteric behavior while enabling tuning of the interaction between aTF and its cognate DNA, including features that arise from sequence architecture.

Sulfane sulfur species, including persulfides and polysulfides, have recently emerged as important signalling molecules,^19^ and key biomarkers in serum and gut microbiota samples,^20–23^ as well as environmentally relevant compounds in wastewater.^24^ However, their high reactivity, transient nature, and chemical interconversion make them challenging to quantify using conventional analytical methods, highlighting the need for sensitive and selective detection strategies.^25^ In this context, transcription factors that directly sense sulfane sulfur provide attractive components for the development of programmable *in vitro* biosensors. We therefore focused on the *reactive sulfur species*–responsive regulator SqrR from *Rhodobacter capsulatus*, a transcriptional repressor that binds the *sqr* promoter. In the presence of sulfane sulfur species, these react with two conserved cysteine residues in SqrR to form a tetrasulfide bond that induces a conformational change, leading to dissociation from DNA and derepression of the promoter.^26,27^ In this work, we applied a SELEX-based selection strategy to engineer DNA operator sequences for SqrR, aiming to optimize aTF-DNA interactions for IVT-based regulation. We identify non-natural operators sequences that preserve ligand responsiveness while enhancing binding affinity and reducing transcriptional leakage. Notably, the selected sequences exhibit binding behaviors consistent with cooperative recruitment of multiple SqrR dimers, suggesting that operator architecture can be leveraged to modulate regulatory performance. Incorporation of these operators into ROSALIND-based transcriptional circuits enables cell-free detection of sulfane sulfur species and establishes operator evolution as a programmable approach to tune aTF–DNA interactions and expand the design space of transcription-based, including for systems lacking well-characterized genetic components.

## RESULTS and DISCUSSION

### *In vitro* selection of new operator DNA sequences

The primary objective of this study was to identify novel DNA sequences capable of functioning as operators for the SqrR protein within an *in vitro* transcription assay. Ultimately, this would contribute to the development of ROSALIND-based biosensors for persulfides or other sulfane sulfur-containing molecules that can trigger SqrR.^25,27^ A recent development of a whole-cell transcriptional biosensor based on SqrR^21^ suggests that it could be feasible to develop an *in vitro* sensor taking advantage of the modularity of the ROSALIND platform. However, the need to optimize an *in vitro* transcription assay in the absence of reducing agents and at increasing concentrations of redox-active species poses a challenge to the range of analyte concentration that could be used. Therefore, we thought of optimizing protein-DNA interactions as part of the first step in the design of the SqrR-based ROSALIND sensor.

Our goal was to find sequences with the highest possible affinity for SqrR, enabling complete transcriptional repression at lower protein concentrations, which has been shown to improve the performance of ROSALIND based biosensors.^7,8^ In addition, these sequences should maintain specificity for the reduced, but not the oxidized state of SqrR, thereby preserving the allosteric connection and the regulatory function of the transcription factor. To this end, we designed an *in vitro* selection process (SELEX). In this method, a random pool of 10^15^ sequences is iteratively enriched in those with the highest affinity for the target. It is important to emphasize that the role of the transcription factor in the ROSALIND assay is to remain bound to the template in the absence of the analyte and dissociate upon its presence. Consequently, we incorporated positive selection steps with the reduced state of SqrR and counter-selection steps against the oxidized state of the protein (**Figure 1A**, Materials and Methods). This was done to ideally obtain new operators with high affinity for SqrR in its reduced form while exhibiting negligible affinity for its oxidized form.

**Figure 1.**
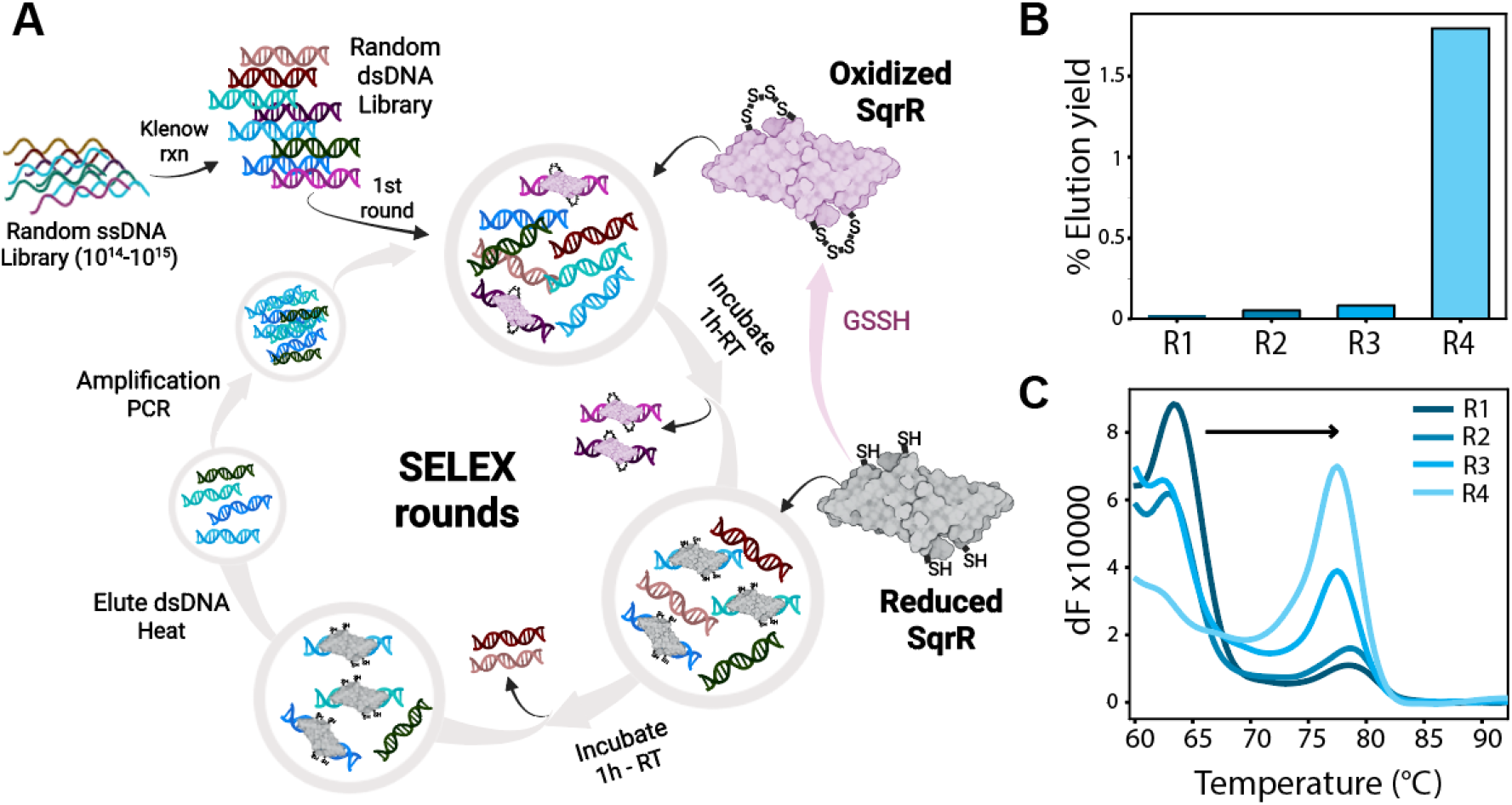
*In vitro* selection of SqrR-binding sequences. **A**. Schematic of the selection workflow. Each round comprised positive selection against reduced SqrR and counter-selection against oxidized SqrR to ensure specificity. **B**. Selection progression was monitored by quantifying elution yields from successive rounds using qPCR. **C**. Melting curve analysis of pools from different rounds was used to evaluate sequence enrichment.

The progress of the selection was monitored using qPCR, determining the elution yield (the ratio of DNA bound to reduced SqrR at the end of a round to the total DNA added at the beginning) and the melting curves of the output DNA, indicative of the diversity of the pools. A significant increase in the elution yield and a notable shift in the melting curves to high temperature were observed by the fourth round (**Figure 1B,C**), indicating the sequence diversity decreases as the affinity of the selected sequences increases. Moreover, to verify the enrichment of high-affinity sequences in the pool, we then turn to evaluate the binding affinity of the initial DNA pool and the pool obtained following Round 4. This assessment was performed via competition assays with the fluorophore labelled natural operator rcc1451 in quantitative fluorescence anisotropy-based DNA-binding experiments. While the oligonucleotides from the initial pool exhibited no measurable competition with rcc1451, the binding curve for the Round 4 pool shifted to the right, indicating that its oligonucleotides effectively compete with the natural operator, with an apparent average affinity one order of magnitude higher than that of rcc1451 (**Figure S1**). Overall, these results indicate that the SELEX process effectively generated an enrichment of the DNA pool with sequences capable of binding to SqrR with affinities comparable to, or even superior to, that of the natural operator.

To identify the individual sequences responsible for this effect, we used high-throughput sequencing (HTS) to analyze sequences from different selection rounds. This enabled us to track the evolution of individual sequences throughout the SELEX process using the FASTAptamer analysis toolkit.^28^ The abundance of unique sequences changed markedly over the four rounds (**Figure S2**), which was consistent with the observed changes in pool diversity from the melting curves. The 500 most abundant sequences (approximately 0.01% of the total unique sequences present in the pool) accounted for roughly 30% of the total reads, demonstrating the highly dominant nature of these high-frequency sequences within the enriched pool.

Taking a closer look at the top 100 most abundant sequences from the last round (Round 4), we performed a motif analysis and generated sequence logos for both the top 100 sequences and the four subpopulations identified in the analysis (**Figure 2A,B**). The first notable observation was a strong enrichment of an ATAT motif among the top sequences. Interestingly, this analysis also recovered and showed a significant enrichment (see Supplementary Text 1) in sequence motifs corresponding to high affinity natural SqrR operators, rcc1451-like (ATTC(N)_8_GAAT) and *sqr*(rcc0785)-like (ATTC(N)_8_ATAT) (**Figure 2C**).^26,27^ However, the selected sequences displayed a pronounced enrichment of ATAT motifs within the (N)□ spacer region, a feature that is not present in the natural operators (**Figure 2A, C**).^26^ The distribution of these motifs within the top 100 sequences reveals a clear enrichment of variants that retain key bases known to be essential for SqrR–operator binding, while simultaneously incorporating novel sequence features that emerged during the selection process. This pattern suggests that the evolved operators preserve critical recognition elements while exploring additional sequence determinants that may modulate the stability and geometry of the SqrR–DNA complex.

**Figure 2.**
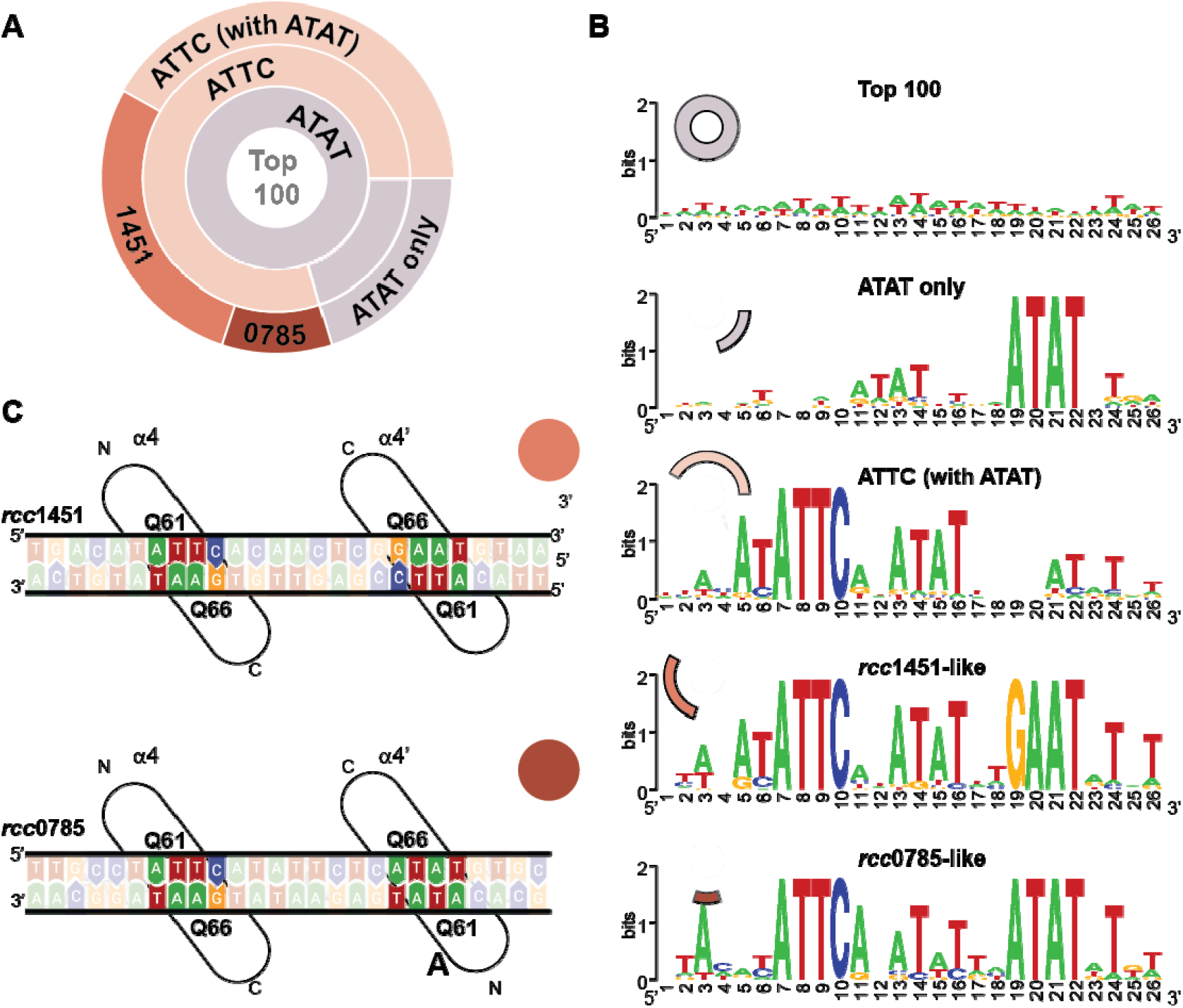
Motif analysis of SqrR-binding sequences. (**A**) Sunburst diagram showing the distribution of sequence motifs within the top 100 most abundant sequences obtained from the SELEX process. The diagram shows the proportion of sequences containing the ATAT motif (100%), those additionally containing the ATTC motif (84%), and those matching the specific patterns of the rcc1451 (ATTC(N)_8_GAAT, 28%) and rcc0785 (ATTC(N)_8_ATAT, 10%) natural operators. (**B**) Sequence logos derived from the top 100 sequences and the four sub-populations identified in the motif analysis. The height of each letter indicates the degree of conservation at that position. (**C**) Schematic representation of the natural SqrR operators rcc1451 and rcc0785. Key residues and bases involved in the protein-DNA contacts are highlighted.

Following the motif analysis, we selected one representative sequence from each of the two major subgroups for detailed individual validation: Seq1, corresponding to the highly dominant rcc1451-like motif, and Seq57 representing the second major subgroup, characterized by the rcc0785-like motif.

### Binding affinity and stoichiometry for the newly selected DNA sequences

To characterize the binding of these two sequences for SqrR, we performed affinity competition assays for both candidate sequences using fluorescence anisotropy. Both 45nt sequences obtained from the SELEX (full sequence) demonstrated a marked increase in binding affinity compared to rcc1451, evidenced by a significant shift to the right in the competition curves (**Figure S3**). In order to avoid potential effects arising from length differences, we turned to truncated versions of Seq1 and Seq57, named Seq1-S and Seq57-S, to match the 26-nt length of the natural operator, rcc1451 (see Material and Methods for details). The results with the truncated sequences confirmed that the affinity was comparable to or higher than rcc1451 (**Figure 3**). While the apparent competition constant for Seq1-S was found to be half that of the reference sequence (Kc = 0.5K), Seq57-S exhibited a significantly higher affinity with Kc = 4K. Beyond the difference in affinity, there was a much more noticeable effect on stoichiometry. In contrast to the 1:1 stoichiometry observed for the natural rcc1451 operator, the competition data for both selected sequences were best described by a multiple-binding site model (**Figure 3A**). Specifically, a model assuming the binding of three SqrR dimers (n = 3) provided an accurate fit across all tested molar excesses for both Seq1-S and Seq57-S (**Figure 3B-C**). These results suggest that the capacity to accommodate multiple SqrR dimers is a conserved structural feature of these enriched sequences, with Seq57-S displaying the highest relative binding strength under the assayed conditions.

**Figure 3.**
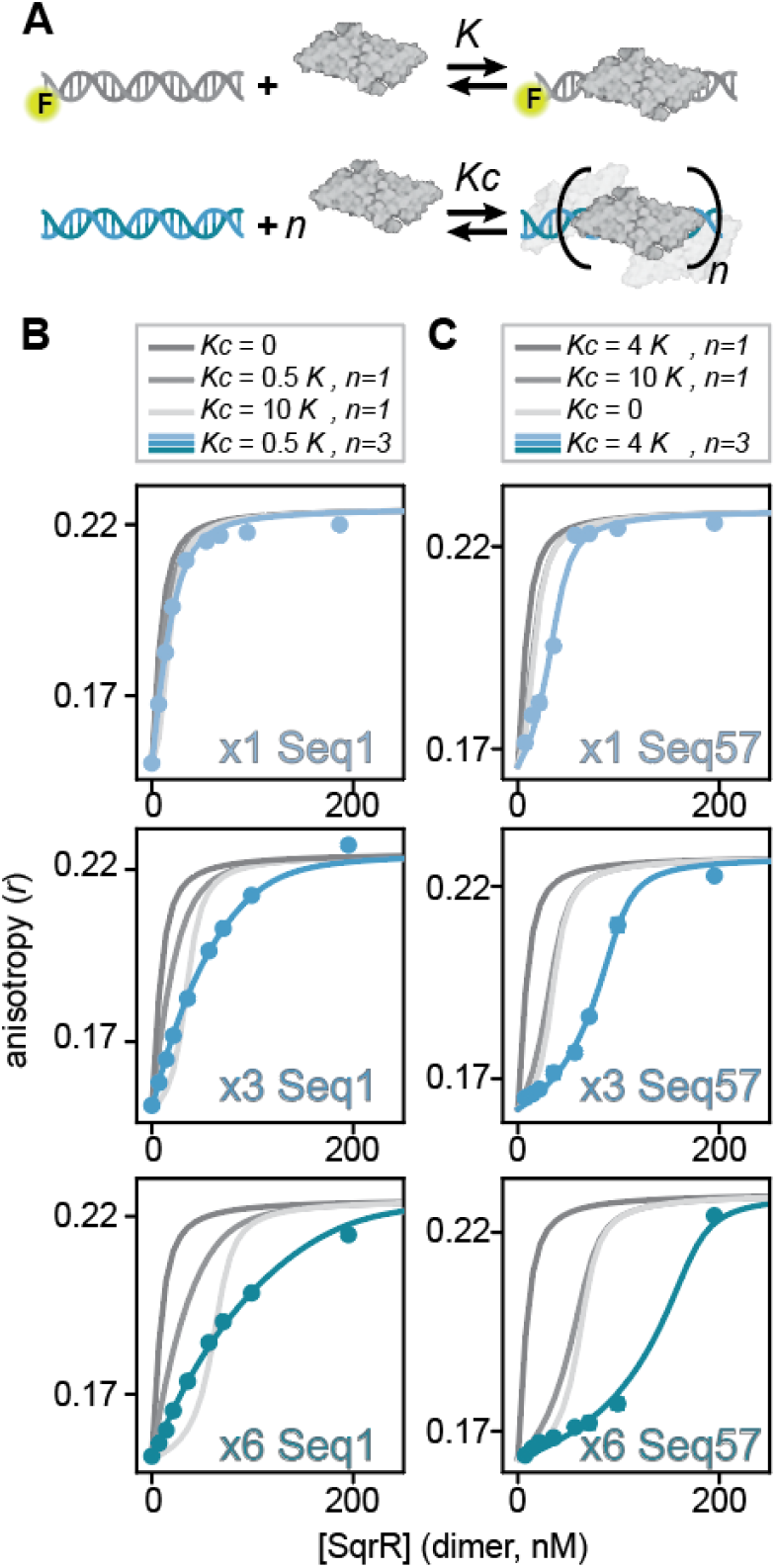
Stoichiometric binding models and anisotropy-based competition assays for Seq1-S and Seq57-S. (**A**) Schematic representation of the proposed binding mechanisms. The natural operator rcc1451 (grey) binds a single SqrR dimer with an affinity constant K. Enriched sequences Seq1-S and Seq57-S (light blue) are modeled to bind n SqrR dimers with a competition constant Kc. (**B-C**) Competition binding curves for (**B**) Seq1-S and (**C**) Seq57-S at 1x, 3x, and 6x molar excesses relative to the reference sequence rcc1451. Experimental data (light blue points) are compared against theoretical models (grey lines) representing various scenarios: no competition (Kc = 0), and models where Kc is a fraction or multiple of K. In both cases, simple 1:1 competition models failed to fit the data. The optimal fit (light blue line) was achieved using a multiple-binding site model with a stoichiometry coefficient of n = 3. For Seq1-S, the fit yielded Kc = 0.5K, while for Seq57-S a higher apparent affinity of Kc = 4K was observed. Best-fit parameters for the simultaneous fitting of the competition binding curves and the direct titration curve of rcc1451 with SqrR are presented in **TableS1** and **FigureS4**.

To validate the multi-valency binding stoichiometry suggested by fluorescence anisotropy, we performed a comprehensive biophysical characterization using Electrophoretic Mobility Shift Assay (EMSA), Size Exclusion Chromatography (SEC), and Mass Photometry (MP). The natural operator rcc1451 consistently exhibited a strictly monomeric binding behavior across all platforms. EMSA experiments revealed a single shifted band regardless of protein excess, with no evidence of higher-order assembly (**Figure 4A**-top). This observation was corroborated by SEC, where the complex eluted as a single, well-defined peak that remained invariant across all tested protein-to-DNA molar ratios (**Figure 4B**-top). Furthermore, MP measurements yielded a molecular mass matching the theoretical value for a single dimer bound to the operator (**Figure 4C**-top). These results align with previous reports for the rcc1451 operator, confirming its function as a canonical binding site that accommodates a single SqrR dimer. In contrast, the SELEX-derived sequences displayed a clear capacity for higher-order recruitment, albeit through distinct assembly mechanisms. For Seq1-S, the binding profile indicated the formation of multiple stoichiometric species. EMSA analysis showed a distribution of discrete, higher-molecular-weight bands as SqrR concentration increased (**Figure 4A**-middle). This heterogeneity was further reflected in the SEC profiles, which exhibited broader peaks shifting toward lower retention volumes (**Figure 4B**-middle), and in the MP data, where a distribution of various complex sizes was observed at a 4-fold protein excess (**Figure 4C**-middle). These findings suggest that for Seq1-S, the assembly of the full multimeric complex involves various intermediate states that coexist under the tested conditions.

**Figure 4.**
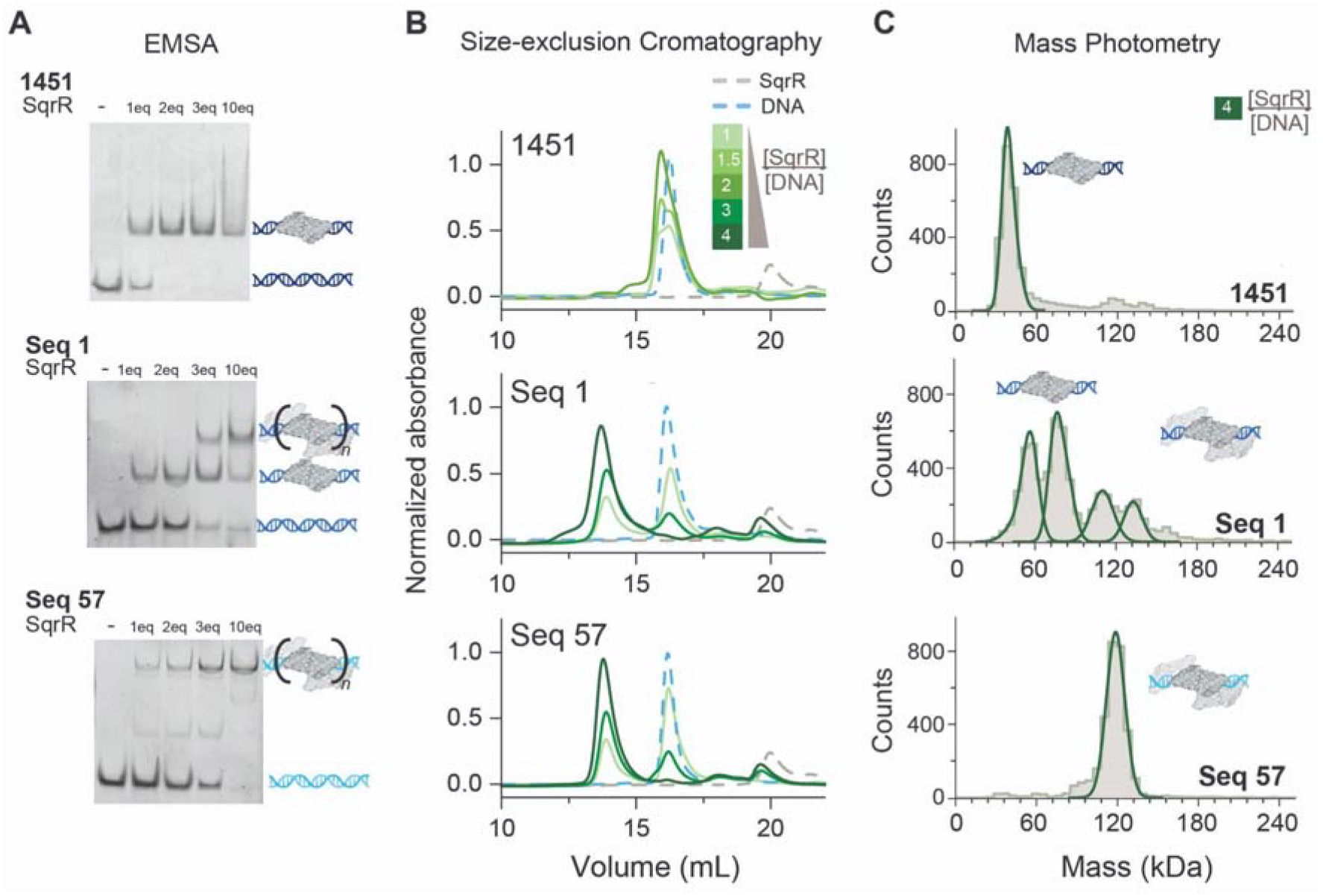
Experimental study of SqrR-DNA binding stoichiometry. (**A**) Electrophoretic Mobility Shift Assay (EMSA) of rcc1451, Seq1-S, and Seq57-S in the presence of 1, 2, 3, and 10 molar equivalents of SqrR dimer. Higher molecular weight bands indicate the formation of multi-dimer complexes for the selected sequences. (**B**) Size Exclusion Chromatography (SEC) elution profiles for SqrR-DNA complexes at various molar ratios. Earlier elution volumes for Seq1-S and Seq57-S complexes relative to the rcc1451 reference indicate the formation of larger macromolecular assemblies. (**C**) Mass Photometry (MP) mass distribution plots for SqrR-DNA complexes at a 4:1 SqrR-dimer:DNA ratio. Measured masses for rcc1451 correspond to a 1:1 stoichiometry, whereas Seq1-S and Seq57-S yield mass peaks consistent with the binding of three SqrR dimers.

However, the most striking behavior was observed for Seq57-S, which exhibited a hallmark mechanism of strong positive cooperativity. Remarkably, EMSA experiments showed the near-exclusive formation of the highest-order multimeric complex, even at stoichiometric ratios (**Figure 4A**-bottom). This ‘all-or-nothing’ binding mode was supported by SEC, which yielded a sharp, narrow peak at low retention times, indicating a highly homogeneous macromolecular assembly. Definitive confirmation was provided by MP, where Seq57-S produced a single, prominent mass peak consistent with the simultaneous recruitment of three SqrR dimers (3:1 stoichiometry).

Overall, the orthogonal evidence from the anisotropy assays, EMSA, SEC, and MP demonstrates that the SELEX process allowed us to identify sequences with high affinity to SqrR, yielding operator sequences that are able to recruit three SqrR dimers, establishing a binding stoichiometry fundamentally different from that of the natural rcc1451 operator. Interestingly, although both Seq1-S and Seq57-S can accommodate three SqrR dimers, the binding profiles indicate different mechanisms, with Seq57-S exhibiting strong positive cooperativity for multi-dimer recruitment, a characteristic not observed for Seq1-S or the natural rcc1451 operator.

### Structural models of selected sequences complexes suggest protein-protein interaction stabilize for trimeric binding

The evidence from the *in vitro* binding characterizations strongly indicates that the selected sequences, Seq1-S and Seq57-S can recruit multiple SqrR dimers. This finding fundamentally contrasts with the 1:1 stoichiometry observed for the natural rcc1451 operator. This remarkable observation suggests the presence of additional putative binding sites or a structural reorganization that facilitates the recruitment of multiple SqrR dimers. To elucidate the molecular determinants governing this altered stoichiometry binding, we conducted a series of bioinformatic analyses and molecular modeling studies, focusing our modeling efforts on Seq57-S due to its pronounced positive cooperativity.

We first employed the AlphaFold Server to generate comparative structural models of three SqrR dimers bound to rcc1451 and Seq57-S. Although the rcc1451 sequence experimentally maintains a 1:1 SqrR dimer-to-DNA binding stoichiometry, AlphaFold predicted the potential conformation of a hypothetical trimeric complex, allowing for a direct structural comparison with the experimentally observed trimeric complexes of the Seq57-S sequence. In both cases AlphaFold positioned the three SqrR dimers along the DNA, with one dimer occupying the known core consensus motif and the other two flanking this central site. A key insight from the modeling was the detailed analysis of protein-protein contacts between the adjacent SqrR dimers. The rcc1451 model, which represents a complex not observed *in vitro*, showed a relatively sparse pattern of inter-dimer protein-protein close contacts. In contrast, the Seq57-S model exhibited a markedly higher number of close contacts between the central SqrR dimer (bound to the known motif) and the flanking dimers (**Figure 5A,B**). These extensive inter-dimer protein-protein contacts in the Seq57-S complex may provide a compelling structural explanation for the formation and stability of the higher-order complexes, consistent with a model of cooperative binding where protein–protein interactions stabilize the overall architecture on the DNA.^11,29^

**Figure 5.**
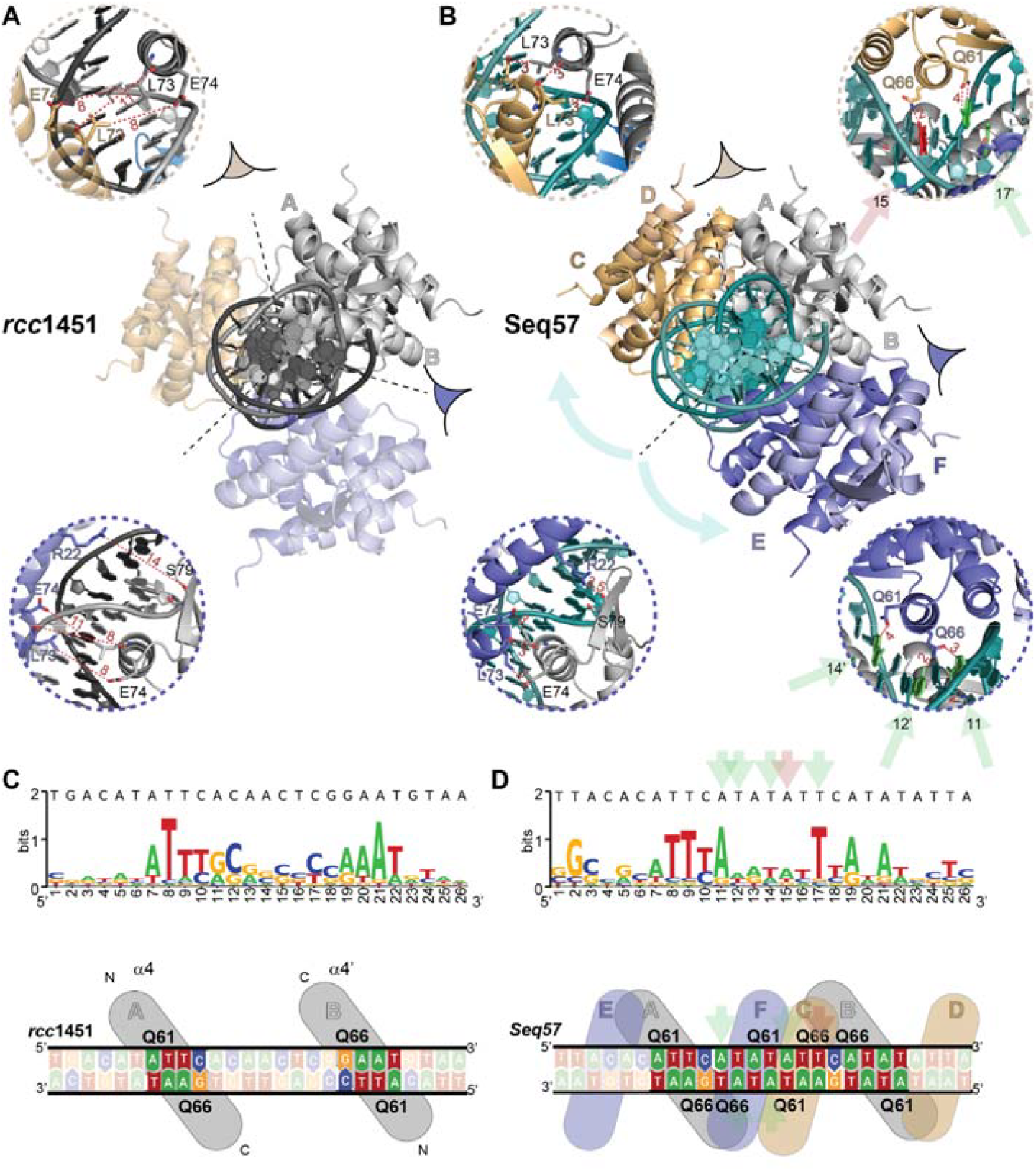
Computational comparison of two SqrR□–DNA complexes. Arrangement of three SqrR dimers bound to the rcc1451 (**A**) and Seq57 (**B**) operators, predicted by AlphaFold3. Insets show magnified views of the interdimer interfaces in both complexes, highlighting the shortest distances detected between residues belonging to different dimers. Distances between residues in helix α4 contacting bases at alternative non-canonical binding sites within the Seq57 operator are also indicated (flanking dimers colored in gold and purple). Sequence logos representing the predicted binding sites corresponding to the rcc1451-SqrR (C, top) and Seq57-(SqrR)_3_ (**D, top**) complexes, in agreement with the experimentally observed stoichiometry. The full operator sequence is shown above each logo for comparison. Schematic representation of the positioning of α4 helices and their residues Q61 and Q66 in base-specific interactions with the rcc1451 (**C, bottom**) and Seq57 (**D, bottom**) operator sequences.

To identify the sequence-level basis for these structural differences, we integrated our structural predictions with sequence-based analysis. Initially, we employed the DeepPBS^30^ geometric deep learning framework to assess how sequence variations affect the binding landscape of AlphaFold-predicted models. This analysis revealed a distinct pattern in the sequence preferences: while the natural rcc1451 operator bound to one SqrR dimer is characterized by a high GC-content at the core, the SELEX-derived Seq57 multidimeric SqrR model shows a marked shift toward a high AT-content (**Figure 5C,D**). This observation perfectly aligns with the motif analysis performed on the most enriched SELEX sequences, where a highly conserved ATAT region is present across the selected operators, while notably absent in the natural rcc1451 (**Figure 2B**). Moreover, in the Seq57 multimeric model, this central AT-rich core facilitates key intermolecular contacts with the flanking SqrR dimers (**Figure 5B-D**). To test the hypothesis that this specific motif is determinant for multimerization, we performed *in silico* mutagenesis on the rcc1451 sequence. Remarkably, introducing only two mutations to generate the ATAT motif in rcc1451 caused the AlphaFold-predicted model to transition from a sparse contact pattern to one nearly identical to Seq57-S, with the dimers positioned significantly closer to facilitate extensive protein-protein interfaces. We further validated this finding experimentally using EMSA. While the wild-type rcc1451 sequence fails to form higher-order complexes even at high protein excess (up to 10 equivalents), the introduction of the ATAT motif was sufficient to induce the formation of the multimeric complex, effectively replicating the stoichiometry of the SELEX-derived sequences (**Figure S5**). These results suggest that the ATAT motif does not merely function as a binding site but may also confer the necessary structural scaffold to optimize the geometry of the protein-protein interface. These findings confirm that the SELEX process that specifically favored the conservation of this central motif that can stabilize the formation of multimeric SqrR-DNA complexes, likely through a protein-protein interface-driven mechanism.

Beyond the static architecture predicted by AlphaFold, the conserved ATAT motif likely confers increased intrinsic flexibility to the selected DNA sequences, consistent with the well-established high flexibility of TpA dinucleotides.^31,32^ Notably, the presence of even a single TpA step has been shown to impact the overall flexibility of DNA molecules.^33,34^ This enhanced flexibility could then enable a favorable conformational change of the DNA helix, better accommodating the three SqrR dimers. Such a conformation could effectively optimize the geometry to maximize the stabilizing protein-protein close contacts, ultimately favoring the formation of the observed stable multimeric SqrR-DNA complexes, a feature that the natural rcc1451 sequence is unable to achieve.

Collectively, the computational data validate the different binding stoichiometries through comparative inter-dimer contact analysis and protein-DNA binding analysis. The results propose that Seq57-S was selected by SELEX based on both a structural scaffold (via new protein-protein contacts) and an enhanced flexibility (via the ATAT motif), promoting highly cooperative multiple binding events (positive cooperativity).

### Functional validation of multivalent operators in cell-free transcriptional biosensors of sulfane-sulfur species

The limited number of reported approaches for sulfane sulfur detection primarily rely on the reactive sulfur species–responsive transcriptional regulators, such as SqrR in whole-cell systems^21,35,36^. To assess its suitability in a cell-free context, we implemented SqrR within a ROSALIND-based platform and evaluated its performance for the *in vitro* detection of sulfane sulfur species. We designed ROSALIND DNA templates incorporating different operator sequences downstream of the T7 promoter (**Table S2**) including natural operators and the multivalent selected operators. The system (**Figure 6A**) relies on the allosteric response of SqrR, which, upon oxidation by sulfane-sulfur species such as GSSH, dissociates from the DNA template to allow the transcription of the 3WJdB light-up RNA aptamer. While we show that the IVT reaction is robust and functions in the absence of any reducing agent, we noted that GSSH concentrations exceeding 500 μM lead to reaction poisoning (**Figure S6**), likely due to non-specific interference with the transcriptional machinery, thereby defining the upper operational limit of the assay.

**Figure 6.**
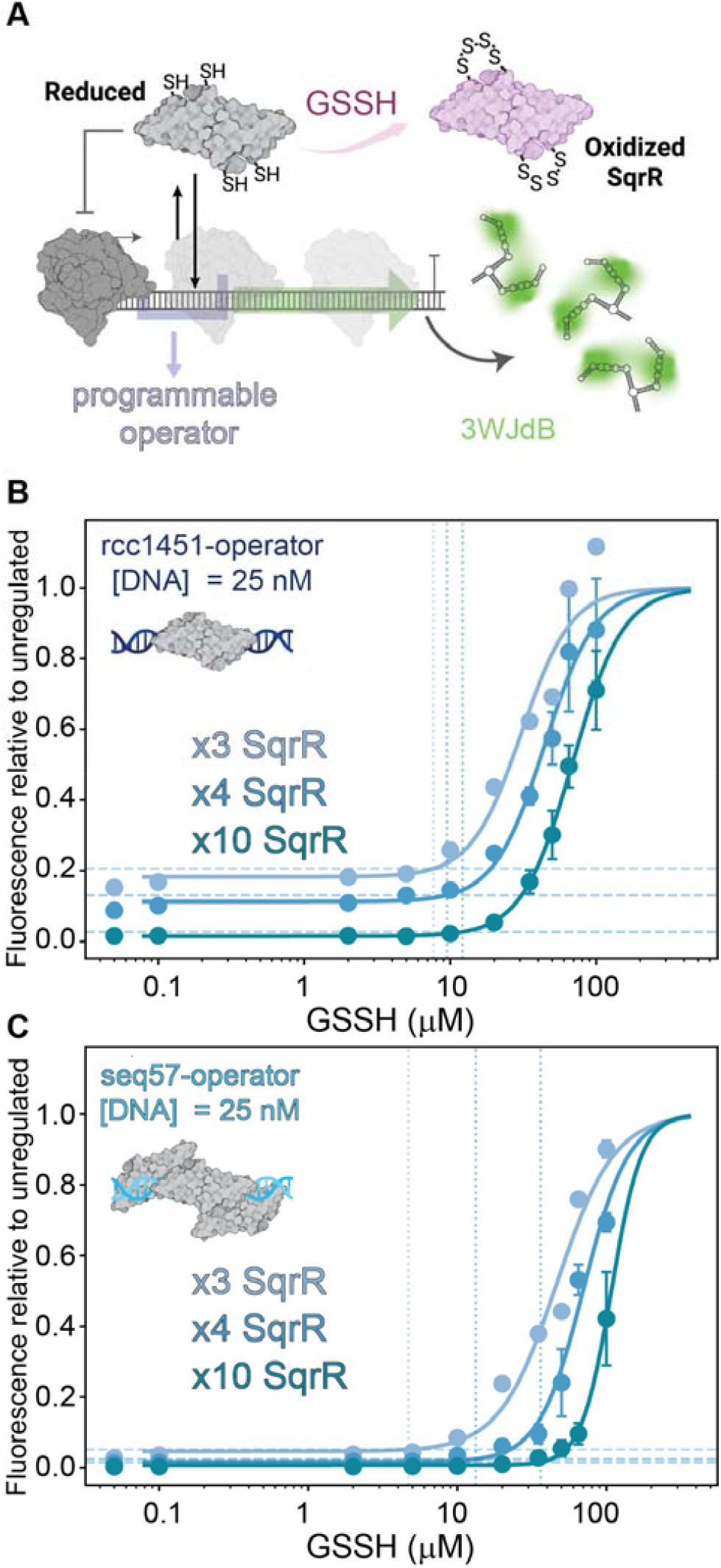
GSSH-induced derepression at different SqrR concentrations of rcc1451 and Seq57. **A**. Scheme of a ROSALIND reaction constructed with SqrR and programmable operator sequences. In presence of sulfane-sulfur species, SqrR becomes oxidized releasing from the DNA, turning transcription on and leading to an observable fluorescent signal from 3 way junction dimeric broccoli (3WJdB) light up RNA aptamer binding to DFHBI-1T. In high concentrations of sulfane sulfurs, such as GSSH, poisoning of reaction is observed (Fig S6) **B. C**. Glutathione persulfide (GSSH) induction dose–response curves of ROSALIND reactions comprised of reduced SqrR dimer at 3x, 4x, and 10x molar excesses relative to DNA template containing rcc1451 (B) and Seq57 (C) operators. Endpoint fluorescence values were read at 1 h and are shown normalized to the promoter-specific unregulated maximum. Data comes from n=2 experiments with 3 technical replicates. High-GSSH concentrations (500 and 1000 µM) were excluded from both display and fit because they fall within the GSSH-poisoning regime. Solid lines are four-parameter Hill induction fits.

We first benchmarked the performance of natural operators for initial characterization. We employed the natural operators rcc1451 and rcc0785, of which only rcc0785 could be successfully incorporated in whole-cell biosensors with a reported limit of detection (LoD) of 25 μM for H_2_S^21^. Comparison of their repression titration profiles (**Figure S7**) showed that rcc0785 requires substantially higher SqrR concentrations to achieve full repression in vitro compared with rcc1451, limiting its suitability for sensitive cell-free applications. We therefore focused on rcc1451 and performed glutathione persulfide (GSSH) induction dose–response experiments (**Figure 6B**).

The rcc1451 operator revealed a characteristic performance trade-off (**Figure 6B**). At low SqrR concentrations, the limit of detection (LoD) is improved, but at the cost of high transcriptional leakage. This high basal signal is a significant bottleneck for molecular biosensing, as it reduces the signal-to-background ratio and could complicate the integration with visual readouts like fluorescence using a portable illuminator^2^ or lateral flow assays (LFA)^8^. Conversely, increasing SqrR concentration effectively suppresses leakage but shifts the LoD toward higher analyte concentrations.

Given that SELEX-derived operators enable multi-dimer recruitment of SqrR, we next asked whether this altered binding architecture translates into improved biosensor performance. We therefore evaluated Seq1 and Seq57 within the ROSALIND platform. Both sequences displayed repression profiles comparable to rcc1451 (**Figure S7**), although Seq1 required slightly higher SqrR concentrations to achieve full repression. Based on its pronounced cooperative binding behavior, we subsequently focused on Seq57 to evaluate the impact of operator-mediated cooperative recruitment on IVT circuit performance. Remarkably, the Seq57-based sensor (**Figure 6C**) showed a nearly absent leakage across all tested SqrR concentrations. This robust repression allows the system to operate at lower protein titers without compromising the baseline, ultimately enabling an improvement in the LoD compared to the natural operator. It is important to note that the normalized fluorescence values reported here correspond to robust, easily detectable absolute signals (**Figure S8**).

The striking difference in behavior between rcc1451 and Seq57 underscores how subtle changes in operator sequence and binding stoichiometry can be leveraged to program distinct sensor performances. Our results demonstrate that SELEX is a powerful tool for fine-tuning the operational window of cell-free circuits, beyond what is available in nature. Ultimately, the successful implementation of this ROSALIND sensor provides a novel and programmable method for the *in vitro* detection of sulfane-sulfur species.

## Conclusions

In this work, we establish an *in vitro* selection strategy to engineer DNA operator sequences that enable tunable and non-canonical transcription factor–DNA interactions in cell-free transcriptional systems. Using SqrR as a model, we show that SELEX can yield operator architectures that preserve ligand-responsive allostery while enhancing binding affinity, reducing transcriptional leakage, and enabling multivalent protein–DNA interactions. These features translate directly into improved performance in IVT-based regulatory circuits, as demonstrated by the implementation of a ROSALIND platform for the *in vitro* detection of sulfane sulfur species. Notably, the identification of operator sequences that promote cooperative recruitment of multiple transcription factor dimers highlights the importance of binding site architecture, beyond affinity alone, in shaping regulatory outcomes.

More broadly, this work provides a generalizable framework for programming transcription factor–DNA interactions through operator evolution, without requiring prior knowledge of native regulatory elements. This approach expands the design space of transcription-based biosensors and may facilitate the development of cell-free systems targeting analytes for which well-characterized genetic components are not available.

