## Supplementary material for "In vitro evolution of DNA operators enables multivalency protein−DNA interactions: towards programmable transcription factor regulation"

**Index**

1. **Material and Methods**
2. **Supplementary Tables**

**Table S1.** Best-fit parameters for the simultaneous fitting of the competition binding curves

**Table S2.** DNA templates used in ROSALIND reactions.

**3. Supplementary text**

**Text S1**

**4. Supplementary Figures**

**Figure S1.** Fluorescence anisotropy-based competition assays for the initial DNA pool and the Round 4 enriched pool.

**Figure S2.** Sequence diversity and enrichment trajectories across selection rounds.

**Figure S3.** Fluorescence anisotropy-based competition assays for Seq1 and Seq57

**Figure S4.** Fluorescence anisotropy assay. Direct titration of the fluorecein-labeled natural operator *rcc1451* with SqrR

**Figure S5.** Introduction of the central ATAT motif in rcc1451 promotes recruitment of multiple SqrR dimers.

**Figure S6.** GSSH poisoning of the unregulated reactions.

**Figure S7.** ROSALIND repression titration curves of natural and selected operators.

**Figure S8**. Time course of unregulated reporter expression.

**1. MATERIAL AND METHODS**

**DNA Library**

All DNA sequences were purchased as synthetic oligonucleotides from Integrated DNA Technologies (IDT). The ssDNA pool acquired consisted of a central random region of 45 nucleotides flanked by two constant sequences at the 3′ and 5′ ends that act as primer regions for amplification (see Table 1). Forward and reverse primers (FwdP and RevP) were used for PCR amplification during the rounds of the in vitro selection, for HTS library preparation, and for qPCR quantification.

The dsDNA library preparation was achieved by conversion of the ssDNA pool into dsDNA by the Klenow reaction. Four individual 25-µL Klenow elongation reactions were set up as follows: 4 µl 100-µM single-stranded oligonucleotide template, 8 µl 100-µM reverse primer, 2 µl 2.5-mM dNTP, 2.5 µl 10×Klenow buffer and 6.5 µl sterile ddH2O. The reaction was heated at 95°C to induce denaturalization of oligonucleotides for 5 min in a PCR thermocycler and then cooled to 25°C at a speed of 0.1°C/s. The mixture was kept at 25°C for 10 min and then 2 µL of the DNA Polymerase I, Large (Klenow) Fragment (New England Biolabs) were added. The reaction was incubated at 25°C for two hours, and then stopped by the addition of 2.78 µL of 100-mM EDTA and heat at 75°C for 20 min. The products were purified through 2% agarose gel and quantified with NanoDrop One (Thermo Fisher).

**Protein preparation**

Proteins were expressed in *Escherichia coli* BL21(DE3)/pLysS cells transformed with a previously reported pSUMO-(C9S)SqrR construct.^1,2^ Cultures were grown at 37°C until OD₆₀₀ = 0.6. Protein expression was induced with 1mM IPTG and cultures were incubated at 30°C for 5h. Cells were collected by centrifugation and resuspended in 40 mL of buffer A composed of 20 mM Tris (pH=8.0), NaCl 500 mM, Imidazole 5 mM, 10% glycerol and 1 mM TCEP, and lysed by sonication at ~4°C. Cellular debris was removed by centrifugation at 15000g. Then, the lysate was further clarified by precipitating nucleic acids with the addition of 0,02% Polyethylenimine (PEI) followed by centrifugation. Proteins were purified using a 10 mL HisTrap column and a gradient of buffer B (same composition as A, but with imidazole 500mM), with SqrR-SUMO eluting around 20% of buffer B. When required, the SUMO tag was removed by incubating with the ULP1 protease^3^ and 5mM DTT overnight at 5°C. Both SqrR and the SUMO-SqrR chimera (used during the SELEX) were further purified by size-exclusion chromatography with a Sephadex G-75 column in isocratic conditions (25 mM Tris pH 8.0, 0.2 M NaCl, 1 mM TCEP, 5% glycerol, 25 °C). Both constructions eluted as homodimers.

**In vitro selection of new sequences**

A schematic diagram of the in vitro selection process is shown in Figure 1.

Protein Immobilization

For all selection rounds, the target proteins were immobilized by binding their His-Tag to magnetic beads (MB) with an NTA-Cobalt-based ligand (Dynabeads™ His-Tag Isolation and Pulldown, Thermo Fisher Scientific). Briefly, 10 µL of MB were incubated with 20 pmol of protein in a total volume of 30 µL of Binding Buffer [Sodium-Phosphate 50 mM, NaCl 300 mM and Tween-20 0.01% (pH 8)] for 15 minutes with gentle agitation at room temperature. The mixture was then placed on a magnetic rack, and the beads were washed three times with 100 µL of Binding Buffer.

First round of in vitro selection

100 pmol of the dsDNA library were mixed with 20 pmol of reduced SqrR-SUMO bound to MB in a total volume of 100 µL of SELEX buffer [HEPES 25 mM and NaCl 200 mM (pH 7)]. The mixture was incubated for 1 hour at room temperature with gentle agitation. After incubation, the beads were placed on a magnetic rack, and the supernatant containing the unbound DNA sequences was carefully removed and discarded and the beads washed three times with 100 µL of SELEX buffer. To elute the SqrR-specific binding sequences, the beads were resuspended in 30 µL of SELEX buffer and heated to 80°C for 5 minutes. The supernatant containing the eluted sequences was then recovered and the elution process was repeated a second time, yielding a final recovered volume of 60 µL of the selected dsDNA library.
1 µL of the eluted dsDNA was used for quantification by qPCR, with the remaining pool used as a template for amplification by PCR (22 cycles of 30 s at 94°C, 15 s at 52°C, 1 min at 72°C, followed by 10 min at 72°C) carried out in 2 tubes of a total volume of 50 µL using a T-Holmes Taq DNA polymerase (Inbio Highway). The PCR product was purified through 2% agarose gel. 1 µL of the purified product was reserved to quantify the amount of DNA by qPCR and remaining DNA was used for the following round.

Second to fourth rounds

60% of the recovered enriched pools from the previous round were mixed with 1 pmol of oxidized SqrR-SUMO bound to MB in a total volume of 100 µL of SELEX buffer as the counterselection step. The mixture was incubated for 1 hour at room temperature with gentle agitation. After incubation, the beads were placed on a magnetic rack, and the supernatant containing the unbound DNA sequences was carefully recovered and mixed with 10 pmol of reduced SqrR-SUMO bound to MB in a total volume of 100 µL of SELEX buffer as the positive selection step. From here, the protocol is the same as for round 1 for sequence elution and PCR amplification. For the fourth round 3 pmol of oxidized protein were used in the counterselection step.

**In vitro selection monitoring**

qPCR was used to monitor the SELEX process in two ways: (i) testing the enrichment of the pools (elution yield) using an absolute quantification and (ii) assessing sequence diversity of the pools by monitoring the melting curve.^4^ Real-time PCR experiments were conducted with the QuantStudio3 qPCR System (Applied Biosystems) according to the manufacturer’s instructions. All reactions were performed in 10 µL of reaction volumes in 96-well plates for PCR. A standard qPCR mixture contained 5 µL of iTaq Universal SYBR Green Supermix (BioRad), 0.3 µL of 10 µM of each primer, 3.4 µL of H2O, and 1 µL of DNA template. Thermal cycling consisted of an initial denaturation at 95°C for 5 min followed by 40 cycles of denaturation at 95°C for 15 s and annealing and extension at 60°C for 1 min. After these amplification cycles, the melting curve analysis was performed from 60° to 95°C. Threshold cycle (Ct) values were determined by automated threshold analysis.

**HTS of selection rounds**

The output of the different rounds were prepared for HTS analysis on an Illumina NovaSeq600 platform (performed by Azenta Inc.), using the NEBNext® Ultra™ II DNA Library Prep Kit for Illumina (New England Biolabs) for library preparation. Briefly, library preparation involved first performing end-repair of fragmented DNA, followed by adaptor ligation, and PCR amplification to produce the final libraries. This kit incorporated different unique dual indexes that allowed the sample analysis of all rounds on one lane. After purification of the PCR product with Agencourt AMPure XP Beads (Beckman Coulter), quantification of the DNA was carried out using a fluorescence method (Qubit kit dsDNA Broad Range), and approximately equal amounts of each library containing specific indexes were mixed. After a quality control (fragment analyzer of the DNA and Qubit), 100–base pair single-end sequencing was carried out.

After demultiplexing, HTS data were analyzed using the FASTAptamer software.^5^ FASTAptamer-Count allows us to count the number of times each sequence is sampled from a population and then rank and sort the sequences by abundance.

**Anisotropy measurements**

The SqrR affinities for the DNA sequences obtained through SELEX were estimated using fluorescence anisotropy-based competition assays, similar to those described previously.^1^ Briefly, a 26-bp fluorescein-deoxythymine (FdT)-labeled operator DNA fragment derived from the rcc1451 promoter region (5’-FdT-GACATATTCACAACTCGGAATGTAA-3’ and its complement) was competed against the unlabeled DNA sequences (obtained through SELEX) at different molar excesses. Reduced SqrR was titrated to saturation, and the resulting binding data were analyzed in DynaFit^6^ using a competition model that accounts for both binding equilibria. To ensure robust fitting, the affinity for the labeled probe (previously determined via direct titration) was fixed. In addition to the 45nt sequences obtained through SELEX (Seq1 and Seq57), we also estimated the affinity for designed shorter versions (Seq1-S and Seq57-S) of these sequences that matched the length of the labeled operator, in order to rule out artifacts arising from length differences. This design preserved the core recognition motif and the overall structural organization of the natural operator with the 16-base-pair central motif flanked by 6 nucleotides upstream and 4 nucleotides downstream, resulting in the desired 26-nt final length. All experiments were performed in duplicate in 25 mM HEPES (pH 7.0), 0.2 M NaCl, 1 mM EDTA, and 2 mMTCEP, 25°C using a Horiba-FluoroMax-4 spectofluorometer.

**Electrophoretic mobility shift assay**

10% continuous polyacrylamide gels were pre-equilibrated in TBE buffer (89 mM Tris, 89 mM boric acid, 2 mM EDTA) by pre-running for 1 h at 120 V at ~4°C (gel box immersed in ice water). Samples (20 µL) containing 100 nM DNA (rcc1451, Seq1-S, or Seq57-S) and 0–10 molar equivalents of reduced SqrR, prepared in the same buffer used for fluorescence anisotropy and supplemented with 5% glycerol, were loaded onto the gel and electrophoresed for ~2.5 h at 90 V. Gels were stained with GelRed for 2 min and imaged using a G-Box Mini transilluminator (Syngene). Images were minimally processed to reduce background.

**Size exclusion chromatography**

Analytical size-exclusion chromatography (SEC) was performed using a Superdex 200 10/300 GL column (GE Healthcare) equilibrated in 25 mM HEPES, 50 mM NaCl, pH 7.0. The column was operated on an ÄKTA FPLC system (Cytiva) at a flow rate of 0.5 mL min⁻¹ at room temperature. Elution was monitored by absorbance at 280 nm. The column was calibrated using standard proteins of known molecular weight. Data were analyzed using Unicorn software.

**Mass photometry**

All mass photometry measurements were performed on a 2MP mass photometer (Refeyn). SqrR-DNA complex samples were prepared mixing 25 nM of the selected DNA operator (rcc1451, Seq57-S or Seq1-S) and 4 equivalents of SqrR in Buffer MP (25mM HEPES pH=7.0, 50mM NaCl, 1mM TCEP). The complex samples were prepared and stored on ice for at least an hour before the MP experiments and the samples were further diluted 10-fold in Buffer MP immediately before the MP measurements were performed.

Precut six-well silicone gaskets were positioned atop precleaned and poly-L-lysine coated glass coverslips to accommodate six samples per coverslip. The poly-L-lysine coating helps DNA adhere to the glass coverslips. These coverslips were subsequently positioned on the stage of a Two-MP mass photometer instrument. Utilizing the lateral control button within the software, the first well was maneuvered over the objective, and 18 µl of PBS buffer was dispensed into one of the gaskets for MP measurement. Subsequently, 2 µl of the diluted SqrR-DNA sample was added and mixed. Sample binding to the poly-l-lysine coverslip was monitored via a one-minute-long movie that was recorded using the acquisition software AcquireMP (AMP) version 2024 R1.1. Standard proteins, such as β-Amylase (BAM; mass: 56 kDa, 112 kDa, 224 kDa) and Thyroglobulin (TG; mass: 670 kDa), underwent measurement in a similar manner as the samples to establish a calibration curve on the same day. Recorded movies were analyzed using the Discover MP (DMP) 2024 R1.0 software. A linear calibration curve was constructed using BAM and TG movies in the DMP software, associating the proteins’ masses with the Ratiometric contrast values and subsequently applied to the sample proteins to ascertain their molecular mass in kDa.

**Computational modeling**

*In silico* structural comparisons of SqrR–DNA complexes were performed using models generated with AlphaFold3 (CITE), which were benchmarked against an experimentally determined crystal structure of the complex (Antelo et al., 2026). Using the AlphaFold3 server, SqrR–DNA complexes were modeled under different stoichiometries, ranging from one to three SqrR dimers bound to different operator DNA sequences. To identify SqrR binding sites on the DNA, we developed custom PyMOL scripts that detect protein–DNA contacts within 4 Å between heavy atoms, consistent with potential H-bond or electrostatic interactions. This approach allowed us to identify the DNA bases most directly involved in interactions with SqrR. To estimate the relative contribution of operator bases to SqrR recognition, the predicted complexes were further analyzed using Deep Predictor of Binding Sites (DeepPBS), a geometric deep-learning model for identifying binding determinants (CITE). Analyses were performed using both readout mode, enabling the calculation of relative importance scores for residues involved in each complex. In addition, custom proximity-based scripts were used to detect potential protein–protein interactions between SqrR dimers within the modeled assemblies. Structural analyses and molecular visualizations were performed using PyMOL (version, CITE)*.*

**ROSALIND reactions**
Homemade IVT reactions were assembled following conditions adapted from established ROSALIND reaction protocols (Jung et al., 2020; Villarruel Dujovne et al., 2025). Transcription templates were generated by PCR amplification of plasmids based on the pJBL729 design (<https://www.addgene.org/140399/>), using primers underlined in the transcription templates listed in Table S1. Amplified templates were purified and verified for the presence of a single DNA band of the expected size on a 2% Tris–acetate–EDTA (TAE) agarose gel. DNA concentrations were determined using a NanoDrop.

rNTP solutions were prepared from solid stocks and adjusted to pH 7.0. Single-use aliquots of IVT components, including DNA templates, rNTPs, 10xIVT buffer (400 mM Tris-HCl pH 8.0, 80 mM MgCl₂, 100 mM DTT, 200 mM NaCl, and 20 mM spermidine), and aTFs were prepared in advance. Unregulated reactions were assembled by adding the following components, listed at their final concentrations, in order: IVT buffer; 0.02 mM (5Z)-5-[(3,5-difluoro-4-hydroxyphenyl)methylene]-3,5-dihydro-2,3-dimethyl-4H-imidazol-4-one (DFHBI-1T); 11.4 mM Tris-buffered nucleotide triphosphates (2.85 mM each, pH 7.0); 0.015 U thermostable inorganic pyrophosphatase; 25 nM DNA transcription template; and Milli-Q H₂O to a total reaction volume of 9 µL. Regulated IVT reactions additionally included purified SqrR and inducer, GSSH. Mixes were equilibrated at 37 °C for 15 min. Immediately prior to plate reader measurements, 1 µL of 50 µM T7 RNA polymerase was added to each reaction. Reactions were characterized using a Varioskan Lux plate reader. Reaction kinetics were monitored by measuring fluorescence (excitation: 487 nm; emission: 510 nm) corresponding to Three-Way Junction dimeric Broccoli (3WJdB)-activated fluorescence. For each condition (n = 3), the data corresponds to the fluorescence intensity measured at 60 min.

**Preparation of RSSH-containing mixtures and SqrR tetrasulfide formation**

Glutathione persulfide (GSSH) was freshly prepared by mixing a 5-fold molar excess of freshly dissolved Na2S with the corresponding thiol disulfide, RSSR. and incubated anaerobically at 30°C for 30 min in degassed 300 mM sodium phosphate (pH 7.4). The concentration of “sulfane” sulfur in the in situ–generated persulfides was determined using a cold cyanolysis assay as previously described,^7^ and these persulfide mixtures were used without further purification at the indicated final concentrations. The same quantification protocol was performed to corroborate the concentrations of the GSSH dilutions prepared for the ROSALIND assays. While these mixtures contain inorganic polysulfides among the “sulfane” sulfur species, the molar fraction RSSH has been reported to be higher than 0.88 of “sulfane” sulfur species^8^. SqrR oxidation was carried out by mixing with a 20 molar excess of GSSH and incubating for 1 h. The excess GSSH was removed by exchanging to fresh buffer using microcentrifuge concentrators.

**Table S1.** Best-fit parameters for the simultaneous fitting of the competition binding curves for Seq1-S and Seq57-S at 1x, 3x, and 6x molar excesses relative to the reference sequence rcc1451 shown in Figure 3. rcc1451 binding constant (K) was set to 3 x10^8^ M^-1^ based on the results of direct titrations (Figure S4), which were compatible with those observed previously for this operator^9^. Conditions: 25 mM HEPES (pH = 7.0), 0.2 M NaCl, 1 mM TCEP, 25°C.

|  | **Kc [x10^8^ M^-1^]** | **number of replicates** |
| --- | --- | --- |
| **Seq1-S** | 1.3 ± 0.02 | 3 |
| **Seq57-S** | 12.0 ± 0.2 | 3 |

**Table S2. DNA templates used in ROSALIND reactions.**

| **Name** | **Operator Sequence** | **Full Sequence, 5' to 3'** |
| --- | --- | --- |
| **pT7- rcc1451-3WJdB-T7t IVT template** | **sqrO**  **tgacatattcacaactcggaatgtaa** | **GCGGATAACAATTTCACACAGGAAACAGCTATGACCATGATTACGCCAAGCTTGCATGCCTGCAGGTCGACTCTAGATAATACGACTCACTATAGGAGGtgacatattcacaactcggaatgtaaCCCACATACTCTGATGATCCGAGACGGTCGGGTCCAGATATTCGTATCTGTCGAGTAGAGTGTGGGCTCGGATCATTCATGGCAAGAGACGGTCGGGTCCAGATATTCGTATCTGTCGAGTAGAGTGTGGGCTCTTGCCATGTGTATGTGGGTAGCATAACCCCTTGGGGCCTCTAAACGGGTCTTGAGGGGTTTTTTG** |
| **pT7- Seq1-S-3WJdB-T7t IVT template** | **Seq1-S**  **gaatgaattcatatatatgaatatat** | **GCGGATAACAATTTCACACAGGAAACAGCTATGACCATGATTACGCCAAGCTTGCATGCCTGCAGGTCGACTCTAGATAATACGACTCACTATAGGAGGgaatgaattcatatatatgaatatatCCCACATACTCTGATGATCCGAGACGGTCGGGTCCAGATATTCGTATCTGTCGAGTAGAGTGTGGGCTCGGATCATTCATGGCAAGAGACGGTCGGGTCCAGATATTCGTATCTGTCGAGTAGAGTGTGGGCTCTTGCCATGTGTATGTGGGTAGCATAACCCCTTGGGGCCTCTAAACGGGTCTTGAGGGGTTTTTTG** |
| **pT7- Seq57-S-3WJdB-T7t IVT template** | **Seq57-S**  **ttacacattcatatattcatatatta** | **GCGGATAACAATTTCACACAGGAAACAGCTATGACCATGATTACGCCAAGCTTGCATGCCTGCAGGTCGACTCTAGATAATACGACTCACTATAGGAGGttacacattcatatattcatatattaCCCACATACTCTGATGATCCGAGACGGTCGGGTCCAGATATTCGTATCTGTCGAGTAGAGTGTGGGCTCGGATCATTCATGGCAAGAGACGGTCGGGTCCAGATATTCGTATCTGTCGAGTAGAGTGTGGGCTCTTGCCATGTGTATGTGGGTAGCATAACCCCTTGGGGCCTCTAAACGGGTCTTGAGGGGTTTTTTG** |

**Supplementary Text**

**Text S1.** To quantify the significance of this enrichment, we compared the observed frequency of these motifs with the expected random probability of their occurrence. For a random sequence of 45 nucleotides (nt), the theoretical probability of spontaneously finding either one of these highly specific 16-nucleotide motifs is less than 0.5% (assuming equal base distribution). However, the analysis of the top 100 sequences revealed a dramatically different distribution, as shown in the sunburst diagrampie chart (Figure 2B): 28% for the 1451 motif and 10% for the 0185. This massive 56- to 20-fold enrichment (relative to the 0.5% expected) conclusively confirms that the recovered sequences are highly selective for SqrR, validating the efficiency of the selection process.

**Supplementary Figures**

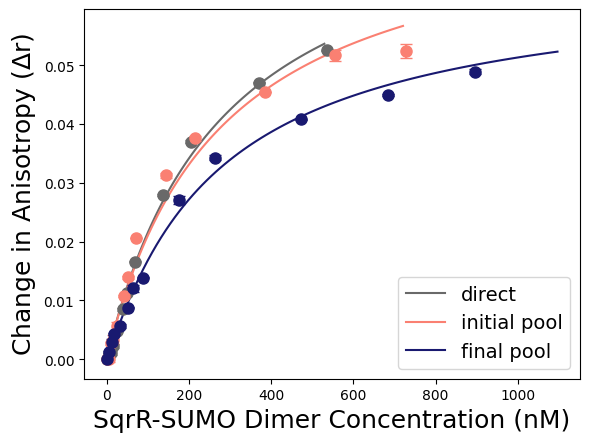

**Figure S1.** Fluorescence anisotropy-based competition assays for the initial DNA pool (orange) and the Round 4 enriched pool (blue). The direct titration curve of the natural operator rcc1451 (grey) is shown as a reference. The oligonucleotides from the initial pool exhibited no measurable competition with rcc1451, consistent with the very low abundance of SqrR-specific sequences at that stage. In contrast, the binding curve for the Round 4 pool shifted to the right, indicating that its oligonucleotides effectively compete with the natural operator. Under these conditions, the apparent average affinity of the Round 4 pool was about one order of magnitude higher than that of rcc1451.

Conditions: 25 mM HEPES (pH = 7.0), 0.2 M NaCl, 1 mM TCEP, 25°C.

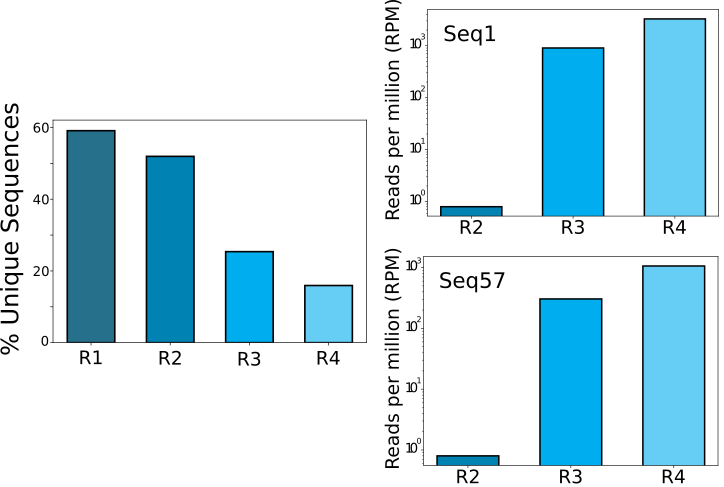

**Figure S2. Sequence diversity and enrichment trajectories across selection rounds.** Left: Percentage of unique sequences identified in each of the four selection rounds. Right: Enrichment trajectories for individual sequences Seq1 and Seq57. Abundance is normalized to Reads Per Million (RPM).

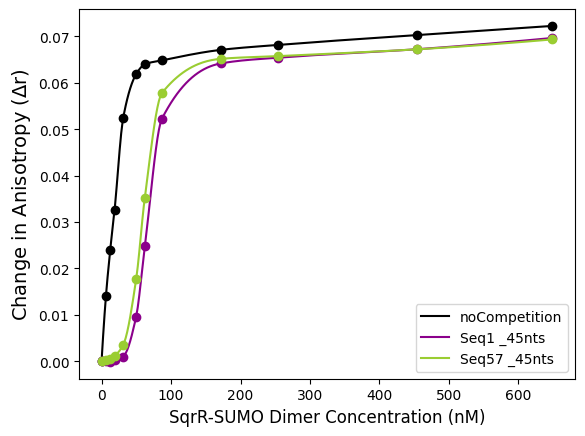
 **Figure S3.** Fluorescence anisotropy-based competition assays for Seq1 (violet) and Seq57 (green). The direct titration curve of the natural operator rcc1451 (black) is shown as a reference. Conditions: 25 mM HEPES (pH = 7.0), 0.2 M NaCl, 1 mM TCEP, 25°C.

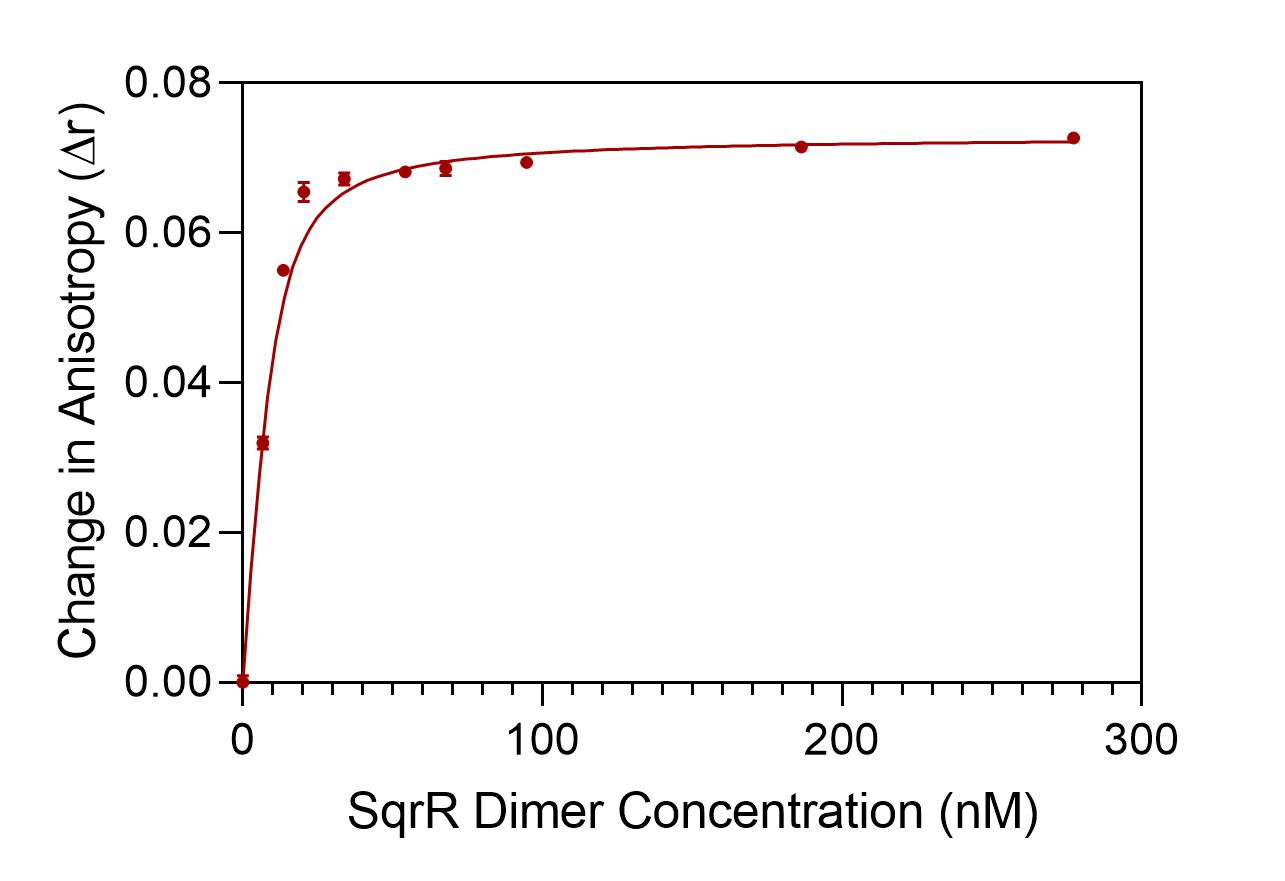

**Figure S4.** Fluorescence anisotropy assay. Direct titration of the fluorecein-labeled natural operator *rcc1451* with SqrR. Data shown correspond to a representative binding isotherm from duplicate experiments. The best-fit binding constant, obtained from a global analysis of all datasets, is (3.0±0.7)×10^8^M^−1^. Conditions: 25 mM HEPES (pH = 7.0), 0.2 M NaCl, 1 mM TCEP, 25°C.

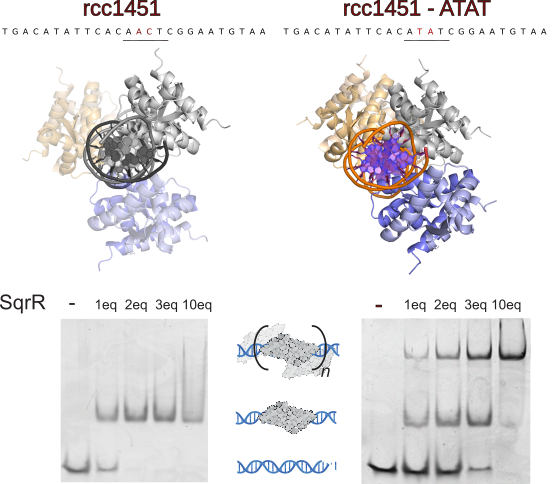

**Figure S5.** **Introduction of the central ATAT motif in rcc1451 promotes recruitment of multiple SqrR dimers**. (Top) Structural models of three SqrR dimers bound to the natural rcc1451 operator (left) and the engineered rcc1451-ATAT mutant (right), as predicted by AlphaFold3. Corresponding DNA sequences are shown above each model; mutated nucleotides in the rcc1451-ATAT variant are highlighted in red to illustrate the introduction of the central ATAT motif. (Bottom) Electrophoretic Mobility Shift Assays for the wild-type and mutated operators in the presence of 1, 2, 3, and 10 molar equivalents of SqrR dimer. Higher molecular weight bands indicate the formation of multi-dimer complexes for the selected sequences.

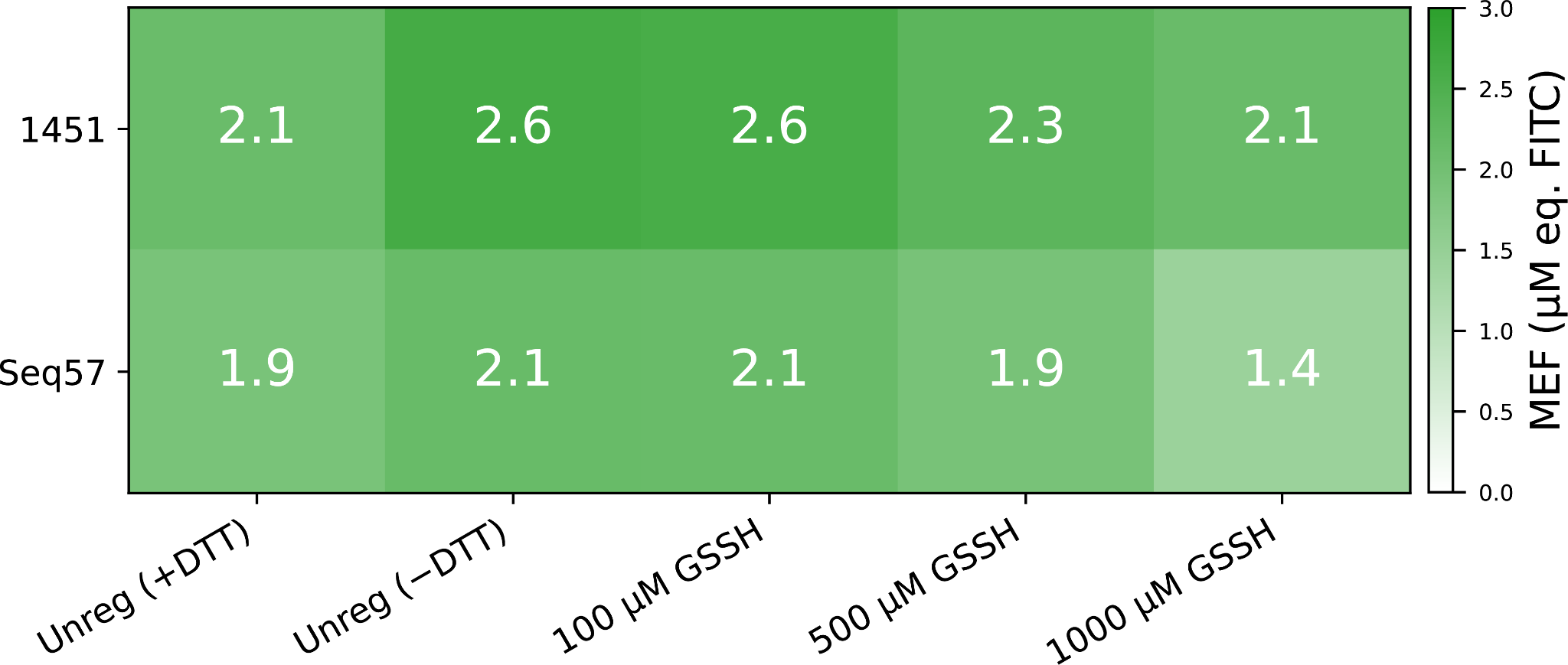

**Figure S6. GSSH poisoning of the unregulated reactions.** The T7 RNAP is typically considered to be sensitive to its redox environment. Thus, IVT and as consequence ROSALIND reactions are usually performed with an excess of a reducing agent such as DTT. ROSALIND reactions have usually 10mM DTT added fresh to their reaction buffer. Here we explored the effect of performing unregulated reactions without any reducing agent and the effect on transcription of sulfane sulfur species such as GSSH. A. A heatmap of endpoint fluorescence from unregulated reactions (lacking *Rc*SqrR) was collected across a GSSH titration (1 h, µM eq. FITC (MEF) units) for two operators (1451, Seq57) across five conditions: unregulated with 10mM DTT, unregulated without DTT (-DTT), and −DTT supplemented with 100, 500 or 1000 µM GSSH. +DTT and −DTT unregulated reactions seem in agreement, indicating that the baseline reducing environment of the ROSALIND mix is already sufficient. In contrast, addition of GSSH to SqrR-free reactions causes a progressive reduction in reporter signal more pronounced at 500 µM. This non-specific effect defines the upper bound of the usable induction window and is the rationale for excluding GSSH concentrations ≥ 500 µM in other experiments.

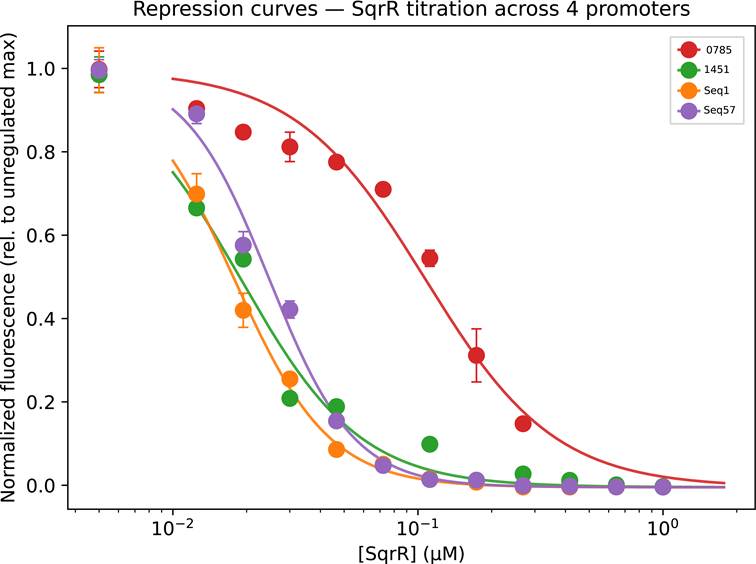

**Figure S7. ROSALIND repression titration curves of natural and selected operators.** Repression dose–response curves with reduced *Rc*SqrR were acquired for four promoter variants, two native-context sequences (0785, 1451) and two SELEX operator designs (Seq1, Seq57). Reduced *Rc*SqrR was titrated across eleven concentrations spanning 0–1 µM (dimer). Endpoint fluorescence was recorded after 1 h of *in vitro* transcription reaction. Data comes from n=2 experiments with 3 technical replicates. Fluorescence is shown normalised to each promoter's unregulated maximum. Solid lines are four-parameter Hill repression fits (y = y_min_ + (y_max_ − y_min_)·K^n^/(K^n^ + x^n^)). The x-axis uses a logarithmic scale with the 0 µM point placed at 0.005 µM for display.

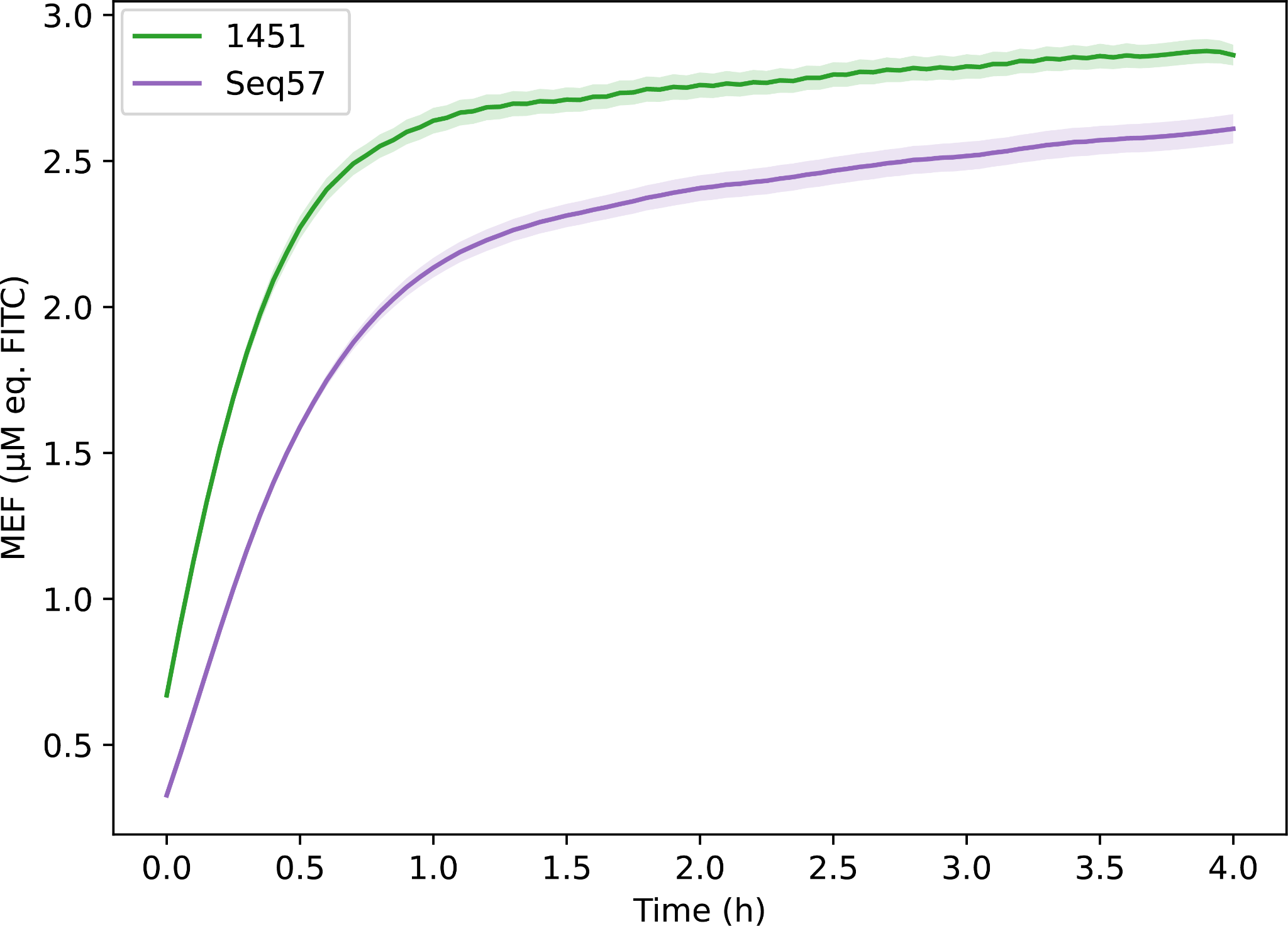

**Figure S8**. **Time course of unregulated reporter expression**. Kinetic traces of the unregulated reactions are shown for rcc1451 and Seq57. Each trace is the per-well mean ± SEM across n=2 experiments with 3 technical replicates with the shaded band denoting the SEM envelope. Fluorescence is reported in MEF units (µM equivalents of FITC).
